# Virulence factors among isolates of extraintestinal *Escherichia coli* (ExPEC) from hospitals in the United Kingdom

**DOI:** 10.1101/2022.10.30.514400

**Authors:** Jane F Turton

## Abstract

182 genes associated with virulence were sought from the whole genome sequences of all non-duplicate isolates of *Escherichia coli* received by the UK Health Security Agency’s Antimicrobial Resistance and HealthCare Associated Infections laboratory for typing between June 2019 and March 2021 from hospitals in the United Kingdom and Republic of Ireland (n=593). These were from healthcare associated investigations and were not associated with diarrhoeal disease. Genes that were very common or very rare were excluded from further analysis. The frequency of detection of genes was compared among isolates from invasive infection, screening, urine samples and from neonates. *cnf1* (coding for cytotoxic necrotising factor), *clbK* (coding for colibactin), *focCDF* (coding for F1 fimbriae), *kpsM*_K1, *neuBD* (associated with the K1 capsule), *mchBC*, *mcmA* (coding for microcins), *papG*_alleleIII (part of a cluster encoding P fimbriae), *pic* (protein involved in intestinal colonization), *sfaE/sfafoCDE* (coding for S fimbriae), *tcpC* (encoding TLR domain containing-protein C) and *vat* (encoding toxin vacuolating autotransporter) were 4 to 28 times more prevalent among isolates from invasive infections than among those from carriage. Representatives of sequence types (ST) 12, 73, 998 and 127 carried multiple of these virulence factors, no matter whether they were from screening swabs or from blood or other infection sites. Isolates carrying multiple virulence factors were more prevalent from neonatal screens than those from general screens. Genes associated with the K1 capsule (*ibeA*, *neuBD*, *kpsM*_K1) were particularly found in STs 1193, 10, 998, 538, 80 and 141. Nanopore sequencing of 14 isolates representing 10 different STs showed that the virulence elements sought were largely carried in chromosomal genomic islands, which were mosaic in nature. Some 9 % of isolates carried more than six of the main virulence genes/gene sets sought, highlighting the potential for significant numbers of carriage isolates to cause extraintestinal infections.

## Introduction

Pathogenic *E. coli* are associated with various infections in humans, the most important being those causing diarrhoea, urinary tract infections, sepsis and meningitis. An extensive array of virulence factors has been described in the organism [1], some of which are characteristic of the disease manifestation. For those causing diarrhoea, various pathotypes have been defined (enteropathogenic *E. coli* (EPEC), enterohemorrhagic (Shiga toxin-producing) *E. coli* (EHEC/STEC), enteroaggregative *E. coli* (EAEC), enterotoxigenic *E. coli* (ETEC), enteroinvasive *E. coli* (EIEC) and diffusely adherent *E. coli* (DAEC), each with their own specific virulence factors e.g. *stx* genes are characteristic of EHEC, production of enterotoxins is characteristic of ETEC and intimin (*eae*) is associated with EPEC [2].

For those isolates not associated with diarrhoeal illness, the situation may not be as clear cut. The patient’s own intestinal flora is the reservoir for these ‘extraintestinal’ pathogenic *E. coli (*ExPEC), which have evolved from commensal strains by acquisition of virulence factors in a cumulative manner [3–6]. Virulence factors described are many and varied and there is often overlap in the virulence gene content among different extraintestinal pathotypes. However, various associations have been made [1]. Uropathogenic (UPEC) *E. coli* are associated with (among others) F1C fimbriae (promoting adhesion), tcpC (TLR domain containing-protein C associated with immune evasion), cytotoxic necrotising factor cnf1, α-haemolysin toxins, pic (protein involved in intestinal colonization), sat (secreted auto transporter toxin) and tsh (temperature sensitive hemagglutinin). Neonatal meningitis-associated (NMEC) *E.coli* are linked with the K1 capsule, with *neuB* and *neuD* genes described as specific for the K1 capsule [7], *ibe* genes (of which *ibeA* is unique to *E. coli* K1), S fimbriae (encoded by the *sfaABCDEFGHS* gene cluster) and also, in common with UPEC, cnf1 toxin; this pathotype can also be responsible for sepsis. Accordingly, *sfaE*, *sfafoCDE* (coding for S fimbriae) and *tcpC* (coding for Toll/interleukin 1 receptor (TIR)-containing proteins) have been associated with sepsis [8]. The toxin vacuolating autotransporter encoded by *vat* has also been significantly associated with bacteraemia [9]. The term NTEC is sometimes used to refer to necrotoxic *E.* coli which express cnf1 [10]. Other potentially important virulence factors include the colibactin biosynthetic cluster in the *pks* pathogenicity island which is often found in combination with *cnf1.* Colibactin represents a class of bacterial genotoxin inducing DNA damage and genomic instability in mammalian cells [11,12]. An analysis of the prevalence of the colibactin island revealed that the *pks* island was consistently associated with the yersiniabactin gene cluster [12,13]. Genotoxic *E. coli* may use colibactin to compete for gut niche utilization [14]. Microcin is also important in establishing colonization in the gut [15,16].

Many of these virulence factors are found as blocks of genes in integrative chromosomal genomic islands referred to as pathogenicity islands (PAI) that are acquired by horizontal transfer [17]. Since they often contain mobile elements derived from bacteriophages, plasmids and insertion sequences, they are subject to rearrangements, insertions and deletions and are therefore variable. These PAI are critical in providing elements encoding functions (such as aiding colonization, immune evasion or adhesin and toxin production) that are essential to the infection process.

In order to understand the importance or otherwise of the various virulence factors described and to be able to identify isolates readily that are of particular virulence among those submitted from healthcare associated investigations (excluding those relating to diarrhoea), we sought 182 virulence factors from whole genome sequences from all isolates submitted to our typing service for *E. coli* from UK hospitals over a period of 20 months during 2019/2020. These largely consisted of those isolated from screens, both for carbapenem resistant organisms and from neonates, but also included clinical isolates from blood, urine, cerebrospinal fluid (CSF) and other sterile sites. While acknowledging that all of the virulence factors play a role, and that gut colonisation is an important step for strains causing extraintestinal infections, we hypothesized that those isolates causing actual infections would be those most likely to possess the full complement of characteristics required to cause those infections. Our panel of isolates therefore provided a context against which to assess the various genes, allowing us to identify a set characterising those of greatest virulence potential.

## Methods

All isolates submitted for cross-infection and outbreak investigation to the UK Health Security Agency’s Antimicrobial Resistance and HealthCare Associated Infection (AMRHAI) laboratory were subjected to whole genome sequencing on a NextSeq Illumina platform following QIASymphony extraction and Nextera® XT library preparation. The bioinformatic pipeline identified the sequence type (ST), predicted O:H type, resistance elements and, where applicable, the ‘SNP address’. The SNP address allows comparison of isolates within the same clonal complex, with groupings at the 0, 5, 10, 25, 50, 100 and 250 SNP levels [18]. The pipeline also sought 182 genes associated with virulence. The set examined, consisting of 593 isolates, were received between June 2019 and March 2021 from 65 hospital trusts from the United Kingdom and Republic of Ireland. These consisted of non-duplicate isolates from CSF (8), blood (53), urine (47), neonatal screens (82), other screens (341) (consisting of ‘CRE screens’, rectal swabs, groin swabs and faeces) and others (ascitic fluid (2), bronchoalveolar lavage (BAL) (4), bile (1), endotracheal aspirate (2), endotracheal tube (ET) secretions (5), endotracheal tube tip (ETT) (5), drain fluid (3), eye (5), fluid (1) foot plantar (1), line tip (1), lung swab (1), lymph node (1), nasopharyngeal aspirate (1), neck (1), pancreatic abscess fluid (1), perinephric collection (1), peritoneal fluid (1), sputum (10) thigh tissue (1), throat (1), wound (5) and unknown (3), and the hospital environment (mascerator (2), cotspace (1), sinkhole (1)). Isolates consisted of 143 different sequence types, of which STs 131, 38, 69, 648, 167, 3168 and 73 were the most common (120, 38, 33, 20, 17, 17 and 14 isolates, respectively). Although the number of isolates from CSF were small (8), they were diverse, with each representing a different sequence type. Phylogenetic groups were identified based on the combinations of the *chuA*, TSPE4C2 and *yjaA* genes [19].

14 isolates were also subjected to nanopore sequencing using a minION Mk1C following DNA extraction with a GeneJet genomic DNA kit (ThermoFisher) and library preparation using the Rapid Barcoding kit SQK-RBK004 (Oxford Nanopore Technologies). Libraries were pooled and concentrated using Agencourt AMPure XP beads (Beckman Coulter). Sequencing was carried out on up to 10 isolates at a time on a FLO-MIN 106D Spot-On flowcell (R9 version) in 72 h runs as recommended by the manufacturer. Virulence genes were further sought using VirulenceFinder 2.0 on the Center for Genomic Epidemiology website (https://cge.cbs.dtu.dk/services/VirulenceFinder/) [20]. BLAST comparisons on a local database were carried out to generate Figure 3 (https://blast.ncbi.nlm.nih.gov/Blast.cgi). Sequences were deposited in the sequence Read Archive under project [PRJNA870093](https://www.ncbi.nlm.nih.gov/bioproject/870093).

## Results

### Highly common genes

Some of the genes were found in all (*aer* (a signal transducer for aerotaxis), *csgABCDEFG* (encoding curli fibers), *ibeC*, *mutS*) or most (*ecpA*, *fimH*, *ibeB*, *matA*) isolates. Others that were relatively common included *chuASTWXY* (associated with haem uptake), *fyuA*, *irp2*, *iucABC*, *iutA* (part of the yersiniabactin and aerobactin siderophore gene clusters).

### Rare genes

Other genes/gene alleles were not found in any of the isolates (*aafABCD*, *aah*, *agg3A*, *agg3B*, *agg3C*, *agg3D*, *agg5A* (encoding aggregative adherence fimbriae), *aidA* (also associated with adherence), *bfpA*, *bmaE*, *cdtA*, *cdtB*, *cdtC* (encoding cytolethal distending toxin), *cofA*, *eae* (encoding intimin associated with ETEC), *eatA*, *espB*, *espC*, *espF*, *etpD*, *f17A*, *fanA*, *fanC*, *fasA*, *fedA*, *fedF*, *fim41A*, *hylE*, *ingA*, *ipaH*, *iucD*, *K88ab*, *itcA*, *neuC*, *nleA*, *papG_*alleleI, *papG_*alleleIprime, *perA*, *pet*, *rfc*, *rpeA*, *saa*, *sta1*, *sta2*, *stb*, *subA*, *tir*). Some others were only found in a very small number of isolates (10 or less) (e.g. *aaiC*, *cif*, *epeA*, *espI*, *espJ*, *iha*, *ipaD*, *ingA*, *katP*, *nleB*, *nleC*, *sepA*, *toxB*, *virF*).

### Main study

As a result of these preliminary observations, the occurrence of the following virulence genes was studied in more detail: *clbK* (part of the colibactin biosynthetic cluster), *cnf1* (encoding cytotoxic necrotising factor 1), *focCDF* (encoding F1 fimbriae), *hylBCD* (encoding α-haemolysin toxins), *ibeA*, *neuBD*, *kpsM_K1* (associated with the K1 capsule), *mchBC*, *mcmA* (encoding microcins), *safE*/*sfafoCDE* (encoding S fimbriae), *tcpC* (encoding TLR domain containing-protein C), *vat* (encoding vacuolating autotransporter toxin), *papG*-_alleleIII, *papC* (encoding pyelonephritis associated pili), *pic* (encoding protein involved in colonization), *tsh* (encoding temperature sensitive hemagglutinin), *sat* (encoding secreted autotransporter toxin) and *malX* (associated with a PAI). The percentage of isolates from various isolation sites carrying these genes is shown in Figure 1 and the distribution of isolates carrying from zero to 8 or more of these virulence factors (where gene clusters are counted as one) is shown in Figure 2. The presence of the rarer *upaH* and *virF* virulence genes was also noted.

**Figure 1.**
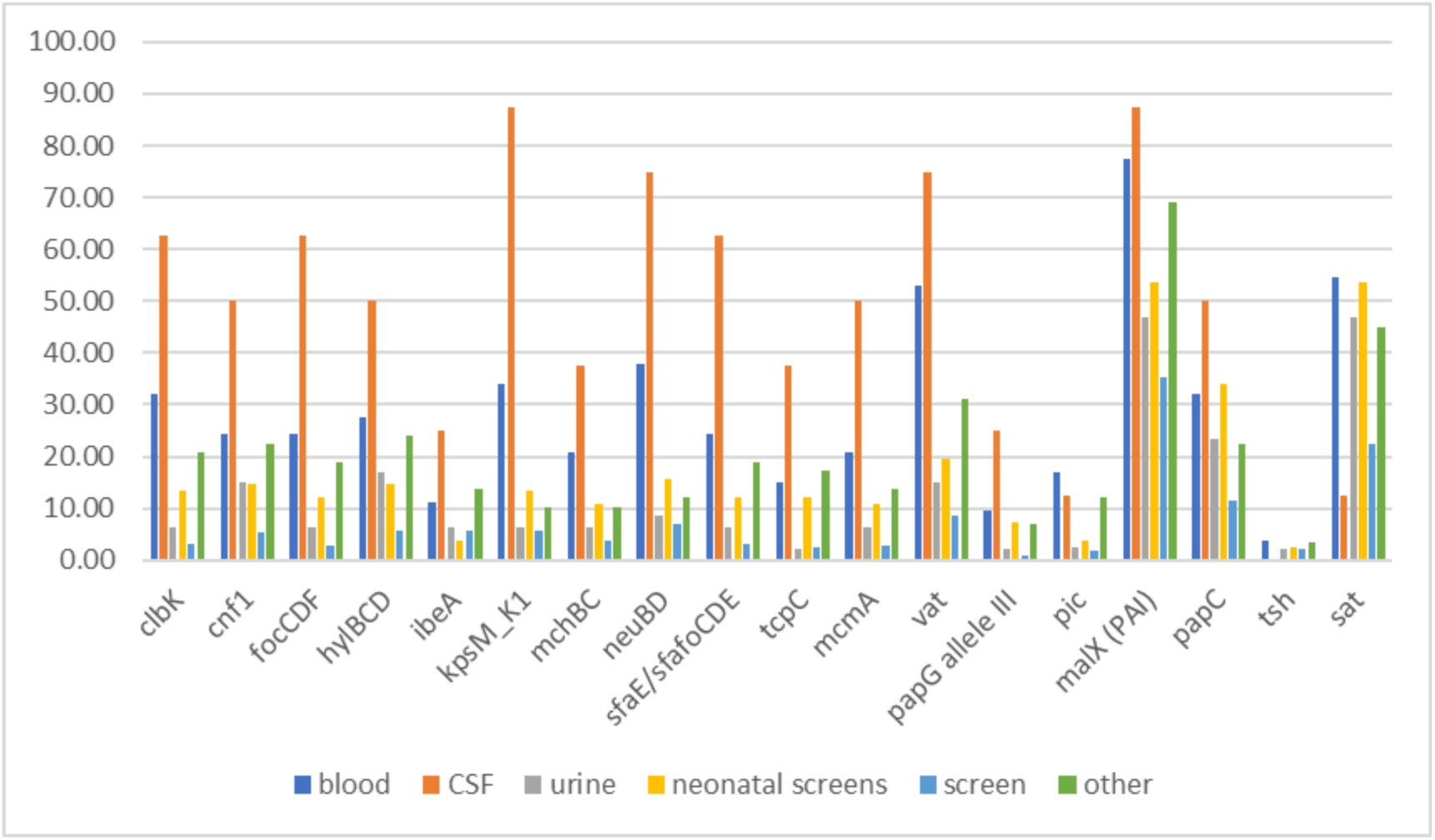
Percentage of isolates from various isolation sites carrying each of 18 virulence genes/gene sets.

**Figure 2.**
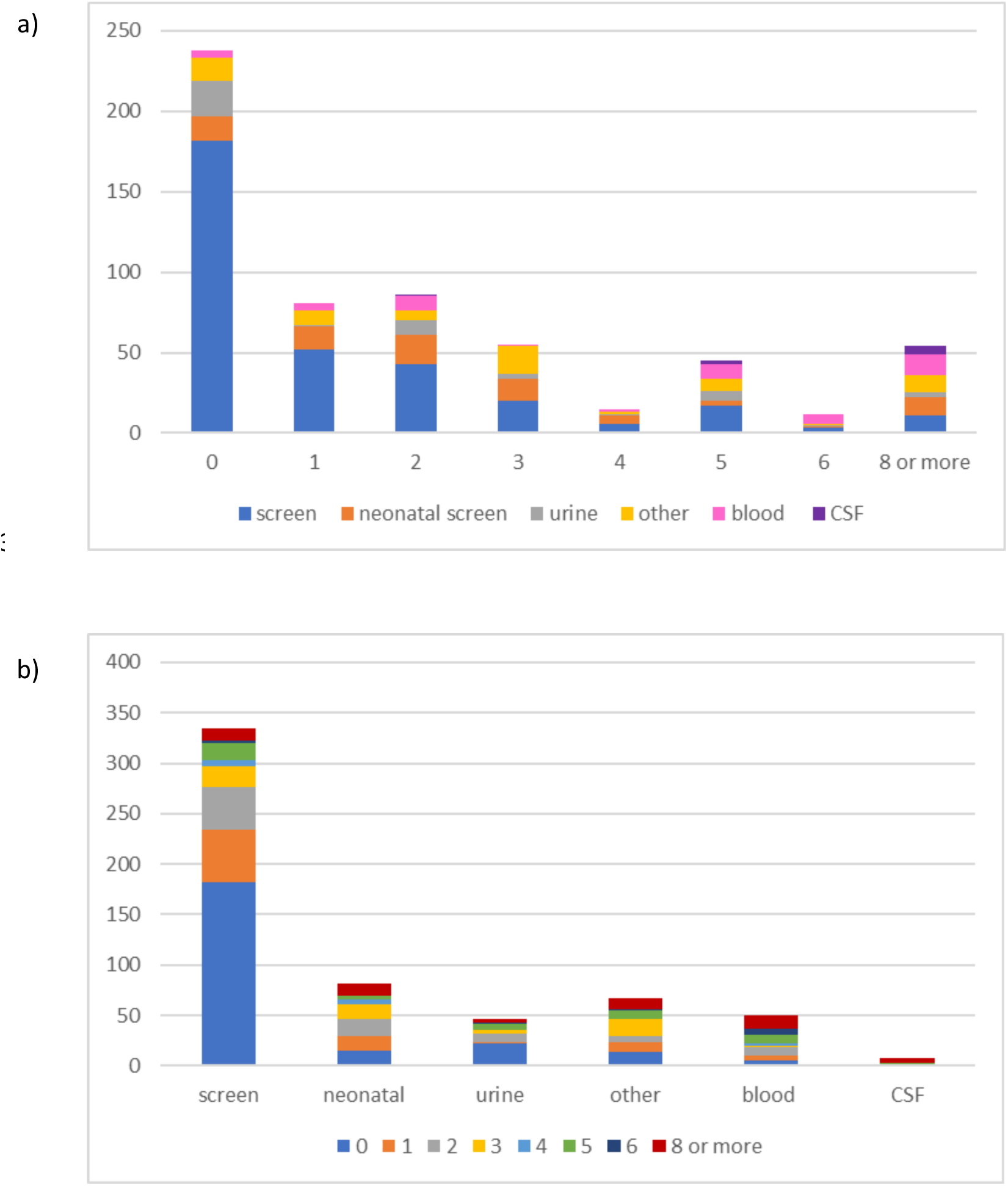
a) Distribution of isolates from different sites carrying 0 to 8 or more virulence genes/gene sets b) Distribution of number of virulence genes/gene sets among isolates from different isolation sites

**Figure 3.**
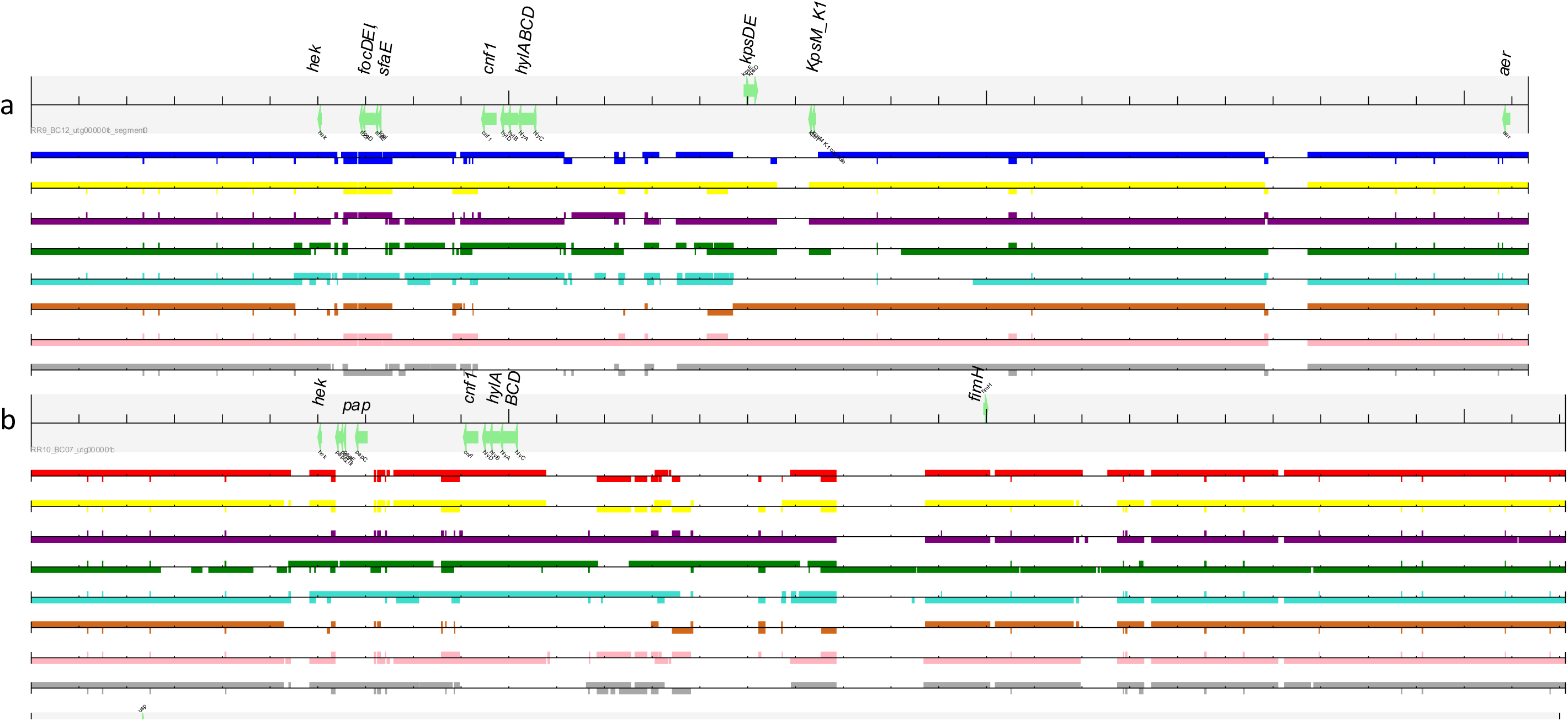
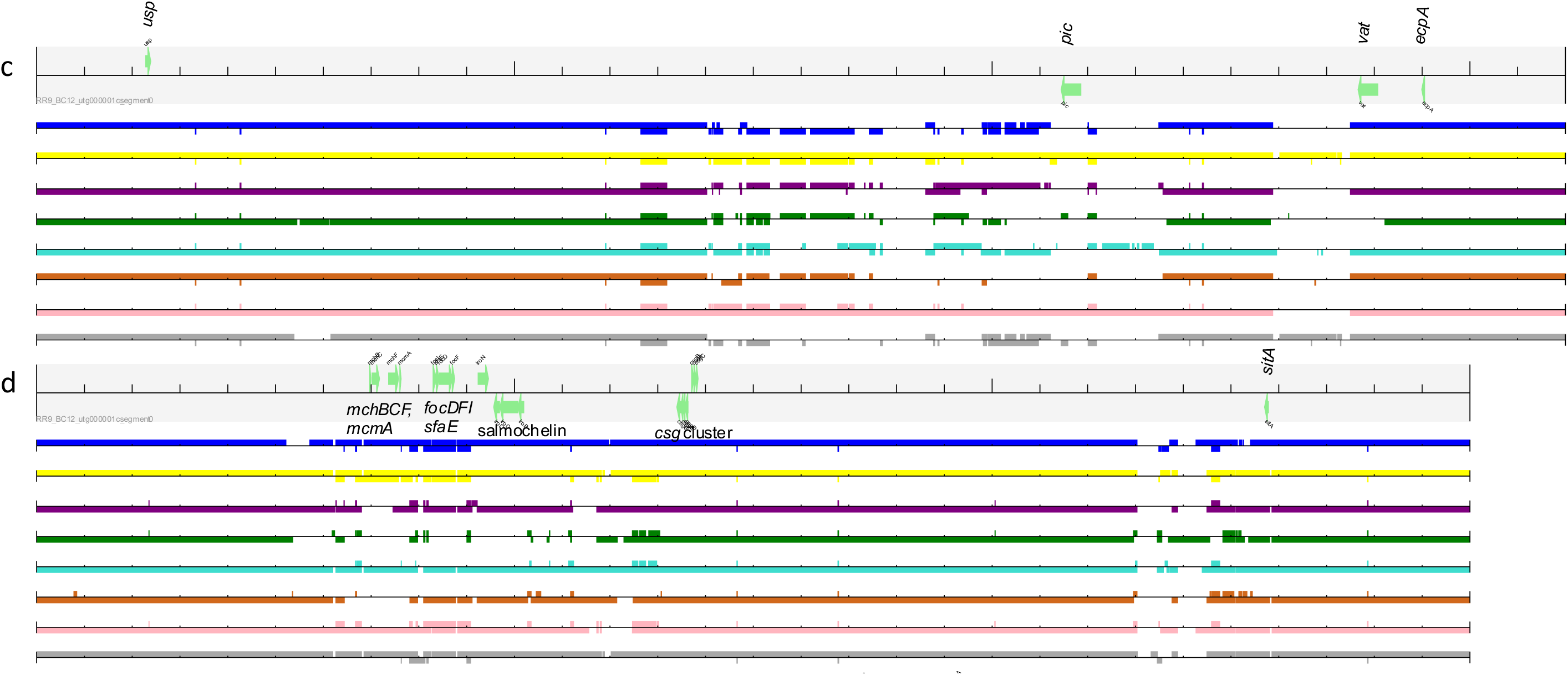
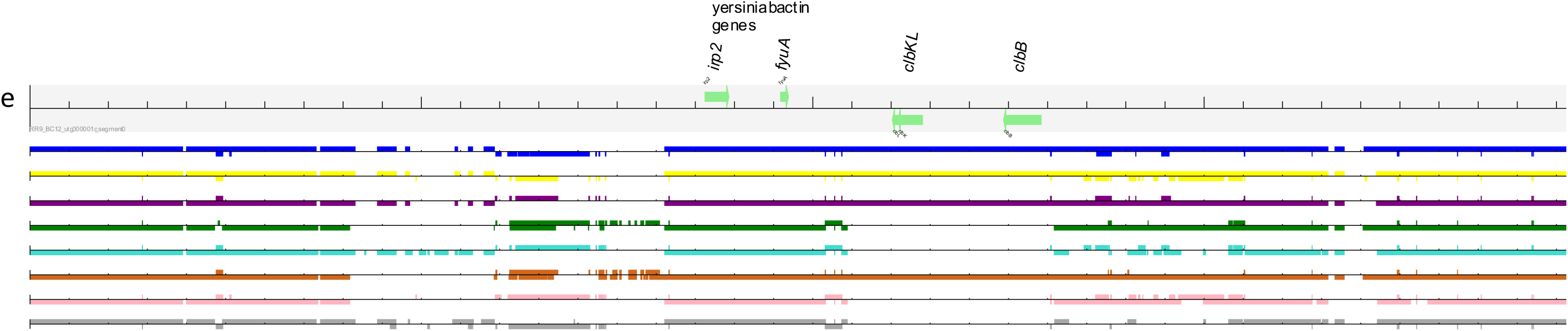
Examples of BLAST comparisons of chromosomal genomic regions carrying multiple virulence genes showing mosaic nature. a) region carrying *focDFI*, *sfaE*, *cnf1*, *hylABCD*, *kpsDET*, *kpsM*_K1 and *aer* genes in P1_SE1_24/20 (ST80; run id RR9_BC12) compared with those in P6_NW1_52/19 (ST12; run id RR10_BC07) shown in dark blue, N12_WM1_31/20 (ST73; run id RR14_BC03) shown in yellow, P11_NE1_45/19 (ST127; run id RR14_BC11) shown in purple, P29_WM3_39/19 (ST131; run id RR14_BC01) shown in green, P12_NW1_38/19 (ST372; run id RR9_BC02) shown in turquoise, N6_S1_40/19 (ST416; run id RR14_BC12) shown in brown, N8_L2_07/20 (ST1859; run id RR9_BC08) shown in pink and N10_L3_26/20 (ST4456; run id RR9_BC07) shown in grey. b) of the same region but comparison against that in P6_NW1_52/19 (ST12; run id RR10_BC07) carrying *papCEFG*, *cnf1*, *hylABCD* and *fimH*; the isolate in red is P1_SE1_24/20 (ST80; run id RR9_BC12) c) region carrying *usp, pic, vat* and *ecpA* in P1_SE1_24/20 (ST80; run id RR9_BC12) compared with that in the other isolates d) region carrying microcin genes (*mchBCF*, *mcmA*), *csg* and salmochelin gene clusters in P1_SE1_24/20 (ST80; run id RR9_BC12) compared with the other isolates e) region carrying yersiniabactin and colibactin in P1_SE1_24/20 (ST80; run id RR9_BC12) Note the mosaic nature of this region carrying multiple virulence genes among the isolates.

### K1 capsule identified by *neuBD* and/or KpsM_K1

The K1 capsule is a PAI encoded protectin that protects the organism from phagocytosis, is associated with bacteraemia and is required for crossing of the blood-brain barrier [17,21].

74/593 isolates were positive for *neuB* (and *neuD*); all but 10 of these were also positive for *KpsM*_K1 indicating that they belong to capsular type K1 associated with neonatal meningitis. 20 (27 %) were from blood, 6 from CSF, 4 from urine, 3 from endotracheal secretions/tube tip, 1 from a diabetic foot lesion, 1 from eye, 1 from sputum and 1 from tissue; the remainder (37) were from screening swabs; at least 24 isolates were from neonates. 10 % of neonatal screens were positive for these elements, while only 7 % of other screens were positive for them. 38 % of blood isolates and 75 % of CSF isolates were positive, suggesting a strong association with sepsis/meningitis. Isolates consisted of 21 different sequence types (STs) of which ST 1193 (23 isolates) and 10 (5 isolates) were the most common, although some of the former were associated with clonal spread among patients related in space and time. No isolates in the entire set were positive for *neuC*.

#### *Ibe* genes

The *ibe* genes are so called because they are associated with invasion of brain endothelial cells. *ibeA* is described as unique to *E. coli* K1, but only some isolates carried both *ibeA* and *neuBD*/KpsM_K1 (19). Every isolate in the entire set (593 isolates) carried *ibeC*; all but one carried *ibeB*. *ibeA* was detected in 39 isolates in total, of which 5 (14 %) were from blood, 2 (5 %) from CSF, 3 from urine, 4 from respiratory samples, 1 from wound and 1 from eye; the remainder (22) were from screening swabs. At least 10 positive isolates were from neonates, but the gene was not more common among isolates from neonatal screens than from screens generally (4 and 6 % respectively). The gene was found in 11 %, 25 % and 6 % respectively of blood, CSF and urine isolates, compared with an overall prevalence of 7 %. The 39 isolates consisted of 22 STs, of which 538 and 998 were the most common, but both were associated with clonal spread among patients linked in space and time.

### S fimbriae (adhesin) genes

Associated with NMEC and UPEC, these are the *sfa* (e.g. *safE*) and *sfafoCDE* genes. The presence of *sfaE* was always accompanied with that of *sfafoCDE*. In total, 54/593 isolates were positive for these genes, of which 5 were from CSF, 13 were from blood, 3 from urine, 1 from tissue, 1 from a line tip, 2 from eye swabs and 7 from other clinical sites; the remainder were from screens (21 isolates) and the environment (1). These genes were found in 25 %, 63 % and 6 % respectively of blood, CSF and urine isolates, compared with a prevalence in general screens of 3 %.

### F1C fimbriae (adhesin) genes

These are associated with UPEC and were found in 53/593 isolates in our panel. We found *focCDF* to be more consistently found than *focG* and *focH*; *focC*, *focD* and *focF* were always found together. Isolates carrying *focCDF* consisted of 5 from CSF, 13 from blood, 3 from urine, 11 from other clinical sites, 10 from neonatal screens, 10 from general screens and 1 from the environment. Prevalence among blood, CSF, urine, neonatal and general screens was 25, 63, 6, 12 and 3 %, respectively, with an overall prevalence of 9 %.

### P fimbriae

P fimbriae are encoded by the *pap* operon (pyelonephritis associated pili) and mediate Gal(α1-4)Gal-specific binding via the adhesin molecule PapG. There are three major alleles of *papG* (GI to GIII) of which alleles II and III are the most common in isolates causing infections in humans [17,23]. We sought *papG*_allele III in particular since we had noted an association with invasive isolates. We detected papG_alleleIII in 22/593 isolates, 6 from blood, 2 from CSF, 1 from thigh tissue, 2 from ET tip, 1 from pus, 1 from urine and the remainder from screens (6 from neonatal screens and 3 from adult rectal screens). The isolates belonged to 12 STs, of which ST12 (9 isolates) was the most common. This allele was found in 9 %, 25 %, 2 %, 7 % and 1 % of isolates from blood, CSF, urine, neonatal screens, and general screens, respectively, compared with an overall prevalence of 3.5 %.

We detected *papC* in 110 isolates (19 %), of which 50 belonged to ST131, 10 to ST69 and 9 to ST12; 17 were from blood, 4 from CSF (representing 50 % of isolates from CSF), 36 from general screens, 28 from neonatal screens, 11 from urine, 13 from other clinical sites and 1 from the environment.

### Cnf-1 (cytotoxic necrotising factor 1)

This exotoxin is associated with NMEC, UTEC and NTEC. It was detected in 68/593 isolates, always, in our experience, with *hylBCD* (encoding α-hemolysin), of which 4 were from CSF, 7 from urine, 13 from blood and 13 from other clinical sites; the remainder were from screens (30) and the environment (1). It was found in 25 %, 50 % and 15 % of blood, CSF and urine isolates respectively compared with an overall prevalence of 11 %. It was detected in 19 STs of which ST131 (16 isolates) and ST73 (12 isolates) were the most common.

### *hylBCD* encoding α-haemolysin

The *hylBCD* gene cluster encodes an exotoxin and is associated with UPEC, including those causing upper *UTIs* such as pyelonephritis; it has been shown to target macrophages [22]. In our study, *hylBCD* were generally found together, with two exceptions only. They were found in 74/593 isolates, of which 8 were from urine, 4 from CSF, 14 from blood and 14 from further clinical sites; the remainder were from neonatal (13), other (20) and environmental (1) screens. The most common ST in which the *hylBCD* cluster was found was ST131 (17 isolates), followed by ST73 (12 isolates).

### TcpC (immune evasion)

This gene encodes an inhibitor homolog of Toll-like receptors (TLR), dampening the proinflammatory response and therefore contributing to immune evasion. It has been associated with both sepsis and UPEC [8]. We detected this in only 41 (out of 593) isolates, only one of which was from urine. The isolates were from CSF (3), blood (8), urine (1), other clinical sites (10) and screens (9 from general screens, 10 from neonatal screens). *tcpC* positive isolates accounted for 15, 38, 2, 12 and 3 % respectively of blood, CSF, urine, neonatal screen and general screen isolates, with a general prevalence of 7 %.

### Colibactin

We sought *clbK*, which is part of the colibactin biosynthetic cluster in the *pks* pathogenicity island. Colibactin is a bacterial genotoxin and is consistently associated with the yersiniabactin (siderophore) gene cluster. It has been associated with persistence in the gut [14]. 60/593 isolates were positive for *clbK*. These consisted of 5 isolates from CSF, 17 from blood, 3 from urine, 1 from tissue, 11 from other clinical sites, 11 from neonatal screens, 11 from general screens and 1 from the environment. These represented 32, 65, 6, 13 and 3 % respectively of isolates from blood, CSF, urine, neonatal screens and other screens, respectively with an overall prevalence of 10 %. Isolates belonged to 19 distinct sequence types, of which STs 73, 12, 1193, 127 and 998 were the most common, in that order. Despite being the most common sequence type (120/593), no representative of ST131 carried *clbk*. *clbK* was detected in all representatives of STs 127 and 998 and in all but one representative of STs 73 and 12.

### Microcins

Siderophore-microcins are antimicrobial peptides that are post-translationally linked to a siderophore which can then kill related bacteria by mimicking iron–siderophore complexes. They therefore provide a competitive advantage in colonization of the gut. They are chromosomally encoded in small genomic islands that, in UPEC, interact with proteins from other genomic islands carrying virulence factors to produce the active siderophore-microcin [16]. We found *mchBC* in only 46 isolates of the set, consisting of isolates from CSF (3), blood (11), line tip (1), pus (1), nasopharyngeal aspirate (1), ET tip (1), sputum (2), urine (3) and screening isolates (9 from neonates, 13 general, 1 environmental) belonging to 22 STs of which ST 73 (12 isolates) and ST12 (8 isolates) were the most common.

#### mcmA (microcin precursors)

Similarly, 46/593 isolates were positive for *mcmA*, and again isolates from CSF (4 isolates), blood (11) and urine (3) were over-represented compared with their overall proportions in the set. Isolates were from 20 different STs, of which ST73 (11 isolates) and ST 12 (8 isolates) were the most common.

### Autotransporters

#### Vacuolating autotransporter toxin (encoded by *vat*)

*vat* was detected in 103 (17 %) of the isolates, 27 from blood, 6 from CSF, 7 from urine, 18 from other clinical sites (ascitic fluid, thigh tissue abscess, pancreatic abscess, ET secretions, line tip, ETT, nasopharyngeal aspirate, sputum, eye, pus), 15 from neonatal screens, 29 from other screens and 1 from the hospital environment (mascerator). These were over-represented in the blood, CSF and other clinical sites groups, representing 53, 75, and 29 % of the isolates in each group, compared with an overall prevalence of 17 % and a prevalence in screens (not neonatal) of only 9 %.

#### Temperature sensitive hemagglutinin (encoded by *tsh*)

The *tsh* gene was found in only 2 % of isolates in our panel. It was not found in any of those from CSF, nor was it found in enhanced numbers in isolates from urine compared to the overall prevalence or from that in screens. It was found in 3.8 % of blood isolates.

#### Secreted autotransporter toxin (encoded by *sat*)

*Sat* was detected in a third (34 %) of the isolates, with most representatives of ST131 (the most common ST by far) carrying the gene. It was found in 55 %, 13 %, 47 %, 54 % and 23 % of blood, CSF, urine, neonatal screens and general screens, respectively.

#### Pic (protein involved in colonization)

The *pic* gene was detected in almost 5 % of isolates overall, with an enhanced prevalence among those from blood (17 %) and CSF (13 %) compared with those from urine and neonatal and general screens (3 %, 4 % and 2 %, respectively).

#### upaH

*u*p*aH* is a further autotransporter gene that has been associated with biofilm formation and bladder colonization [24]. Among our panel of isolates, we only detected it in 18 isolates, all but one of which belonged to ST73 or a single locus variant (SLV) of ST73. This ST generally is associated with virulence; not surprisingly 7 of the 18 isolates were from blood and 2 from line-tips, while only 5 were from screens, despite the latter greatly outnumbering those from blood (53 blood isolates compared with 423 isolates from screening). The isolates were from 10 different hospitals, but 4 of them were clonally related to each other (5-SNP cluster) and were related in space and time.

### PAI marker gene *malX*

*malX* codes for a phosphotransferase system enzyme that recognizes maltose and glucose. It is a PAI marker. It was detected in almost all representatives of ST131 (117/ 120 isolates) and was common among the panel, being found in 277/593 isolates (47 %) overall, with somewhat increased prevalence among blood (41/53) and CSF isolates (7/8) (77 and 89 %, respectively) compared with those from urine, neonatal and general screens (47, 54 and 35% respectively). As well as ST131, it was strongly associated with STs 648 (18 isolates), 1193 (24 isolates) and ST73 (14 isolates).

### Prevalence of virulence genes in invasive isolates compared with screens

These data do not provide sufficient support that *tsh* is more associated with invasive isolates compared with screen isolates. However, there is strong evidence in support of *focCDF*, *sfaE/sfafoCDE*, *clbK*, *papG*_alleleIII, *pic* and *mcmA* being associated with sepsis (all greater than 7 times more prevalent among blood isolates than screen isolates), *focCDF*, *sfaE/sfafoCDE*, *papG*_alleleIII, *cblK* and *mcmA* associated with CSF (all greater than 17 times more prevalent among CSF isolates than general screens and more than 6 times more prevalent among CSF isolates than overall) as well as *cnf1*, *kpsM*_K1, *mchBC*, *neuBD*, *vat* and *tcpC* . *hylBCD* and *cnf1* were associated with urinary tract infections (both greater than 2.8 times as prevalent among urine isolates than overall). A caveat to these observations is that the number of isolates from CSF was small (8/593 isolates); however, they were genetically diverse, each representing a distinct ST.

### Combinations of virulence genes

While some 355 isolates of the panel of 593 carried one or more of the virulence genes (or virulence gene sets) sought (*clbK*, *cnf1*, *focCDF*, *ibeA*, *hylBCD*, *mchBC*, *neuBD*, *ibeA*, *sfaE*/*sfafoCDE*, *tcpC*, *KpsM*_K1, *malX* (PAI), *mcmA*, *papC*, *papG*, *papG*_alleleIII, *pic*, *sat*, *sepA*, *tsh*, *upaH*, *vat*, *virF*), 81 carried only one (particularly *malX*, *sat*, *tsh*, *papC*, or in one instance the rare gene *virF* found in only 2 isolates in our set), 86 carried only two (with the most common combinations being *malX* and *sat* (62) and *papC* and *sat* (8), and 49 carried only 3 (most, all of ST 131, carrying the *malX*, *papC* and *sat* combination (30)). A number of isolates including 12 from invasive sites carried the combinations of *neuBD*, *KpsM*_K1, *malX*, *sat*, *vat* (15 isolates, all of ST 1193), *neuBD*, *KpsM*_K1, *malX*, *papC*, *sat* (5 isolates) or *ibeA*, *neuBD*, *KpsM*_K1, *malX*, *vat* (7 isolates) (with 10 further isolates carrying one of these combinations and further genes), while the combination of *cnf1*, *hylBCD*, *malX*, *papC*, *sat* was found in 15 isolates (all of ST131), of which 4 were from urine (with a further 6 isolates carrying this combination and further genes). Some 53 isolates (8.9 %) carried more than 6 of these genes/gene sets, and all but one of these carried at least 8; one carried 15 (Table S1). Of these 53 isolates, which all belonged to phylogroup B2, 47 carried *cblK*; this represents 78 % of all the isolates carrying *cblK*, showing that isolates carrying this gene are highly likely to carry multiple virulence factors. Many (16) carried one or more elements associated with the K1 capsule (*KpsM*_K1, *ibeA*, *neuBD*). All but three carried the F1 fimbriae and S fimbriae genes *focCDF* and *sfaE*/*sfafoCDE*. Sequence types 73 (13 isolates, plus 2 SLVs), 12 (8 isolates), 127 (6 isolates) and 998 (6 isolates, 4 from neonates from the same hospital and 3 representing the same strain) were the most common STs among those carrying 8 or more virulence genes/gene sets. Worrying, eight also carried *bla*_OXA-48-like_ carbapenemase genes. In common with 3 of the 4 ST998 isolates, and 4 of the ST73 isolates, the two isolates belonging to ST416 and two of those belonging to ST80 were epidemiologically and clonally related; all the others were not. A further two of the 13 ST73 isolates were from the same hospital and submitted together but were more than 250 SNPs apart.

### Nanopore sequencing results

Nanopore sequencing was carried out on 14 isolates, consisting of 10 different sequence types (STs 12, 73, 80, 127, 131, 141, 72, 416, 1859 and 4456 (Table 1). Six of the isolates carried elements associated with the K1 capsule (*KpsM*_K1, *ibeA*, *neuBD*). In most cases (9/14), the assemblies resulted in complete closed circular contigs for the chromosome and any plasmids present; these were those included in comparisons in Figure 3. It confirmed that the virulence genes sought were mostly chromosomally located in these examples. An exception was the 128 kb plasmid in N6_S1_40/19, which carried the aerobactin and salmochelin clusters, *iss*, *ompT*, *sitA* and *traJ* genes. In other isolates, these elements, where detected, (with the exception of *traT* and *traJ*) were chromosomally located.

**Table 1.** Isolates subjected to nanopore sequencing. Each isolate was from a different patient and, with two exceptions only (hospitals NW1 and WM1), from a different hospital. Isolates were labelled by patient (P1-P29, N1-N24, where those beginning with N are from neonates), hospital (by region and number within that region) and date of isolation (week/year). Regions were EE, East of England, EM, East Midlands, WM, West Midlands, L, London, NE, North East, NW, North West, SE, South East, SW, South West, S, Scotland, W, Wales, Y&H, Yorkshire and Humber.

| Isolate | ST | SNP address | O:H type | Isolation site | Carbapen-emase gene | Virulence genes |
| --- | --- | --- | --- | --- | --- | --- |
| P6_NW1_52/19 | 12 | 13.13.13.13.13.13.13 | O4:H5 | rectal screen | *bla<sub>OXA-48</sub>* | *clbK, cnf1, focCDF, hylBCD, mchBC, sfaE, tcpC, malX, mcmA, papC, papG_alleleIII, vat* |
| P7_NW2_18/20 | 12 | 1.16.16.16.16.16.17 | O4:H5 | blood | none detected | *clbK, cnf1, focCDF, hylBCD, mchBC, sfaE, tcpC, malX, mcmA, papC, papG_alleleIII, vat* |
| N9_EE1_25/20 | 12 | 1.17.17.17.17.17.18 | O4:H5 | faeces | none detected | *cnf1, focCDF, hylBCD, mchBC, sfaE, tcpC, malX, mcmA, papC, papG_alleleIII, vat* |
| P11_NE1_45/19 | 127 | 3.9.9.9.19.20.20 | O6:H31 | kidney urine | *bla<sub>OXA-48</sub>* | *clbK, cnf1, focCDF, hylBCD, sfaE, tcpC, malX, mcmA, papC, vat* |
| P29_WM3_39/19 | 131 | 1.4.441.470.476.510.551 | O25:H4 | urine | none detected | *cnf1, hylBCD, malX, papC, sat* |
| P4_EM1_41/19 | 141 |  | O50O2:H6 | CSF | none detected | *clbK, cnf1, focCDF, hylBCD, mchBC, neuBD, sfaE, tcpC, KpsM_K1, malX, mcmA, vat* |
| N8_L2_07/20 | 1859 |  | O99:H6 | blood | none detected | *cnf1, focCDF, hylBCD, ibeA, mchBC, neuBD, sfaE, KpsM_K1, malX, mcmA, pic, vat* |
| P12_NW1_38/19 | 372 |  | O83:H31 | CSU | *bla<sub>OXA-48</sub>* | *cnf1, focCDF, hylBCD, ibeA, mchBC, sfaE, malX, mcmA, papC, papG_alleleIII, vat* |
| N6_S1_40/19 | 416<br>(CC95) | 6.27.28.28.28.29.53 | O18ac:H7 | blood | none detected | *clbK, focCDF, ibeA, neuBD, sfaE, KpsM_K1, malX, vat* |
| N10_L3_26/20 | 4456 |  | O83:H4 | blood culture | none detected | *focCDF, ibeA, mchBC, neuBD, sfaE, KpsM_K1, malX, mcmA, papC, papG, vat* |
| N5_WM1_04/20 | 73 | 56.70.70.71.72.73.74 | O6:H1 | line tip | none detected | *clbK, cnf1, focCDF, hylBCD, mchBC, sfaE, tcpC, malX, mcmA, pic, sat, upaH, vat* |
| P15_Y&H1_29/20 | 73 | 59.74.75.76.77.78.79 | O6:H1 | blood | none detected | *clbK, cnf1, focCDF, hylBCD, mchBC, sfaE, tcpC, malX, mcmA, papC, pic, sat, upaH\*, vat* |
| N12_WM1_31/20 | 73 | 61.76.77.78.79.80.83 | O6:H1 | nasopharyngeal aspirate | none detected | *clbK, cnf1, focCDF, hylBCD, mchBC, sfaE, tcpC, malX, mcmA, pic, upaH, vat* |
| P1_SE1_24/20 | 80 |  | O75:H7 | urine | none detected | *clbK, cnf1, focCDF, hylBCD, ibeA, mchBC, neuBD, sfaE, KpsM_K1, malX, mcmA, pic, vat* |

All the isolates carried *chuASTWXYU* (encoding haem uptake) in the chromosome in a gene cluster that was distant from the other virulence genes sought. The three representatives of ST12 from 3 different hospitals shared a similar arrangement in terms of order of genes detected by the VirulenceFinder software with all sharing a section of approximately 214 kb containing the genes *sitA* (coding for an iron transport protein), *iss* (increased serum survival), the salmochelin cluster (e.g. *iroN* encoding a siderophore protein), *focC*, *sfaD* (coding for S fimbriae/F1C minor subunit), *mcmA*, *mchF*, *mchC*, *mchB* (encoding microcins) and *cea* (coding for colicin E1) (in that order). This region also included the *csgABCDEFG* cluster (coding for curli fibers) and *agn43* (coding for antigen 43 involved in biofilm formation). The 3 isolates shared high overall similarity with one another (with P7_NW2_18/20 having 99.5 % identity, 99 % coverage with N9_EE1_25/20 while P6_NW1_52/19 shared 99.95 % identity and 98 % coverage with it), with all carrying *terC* (encoding tellurium ion resistance protein), *clbB* (coding for hybrid non-ribosomal peptide/polypeptide megasynthase), *yfcV* (encoding a fimbrial protein), *gad* (coding for glutamate decarboxylase), *ompT* (encoding an outer membrane protease), *vat* (encoding vacuolating autotransporter toxin), *usp* (encoding uropathogenic specific protein), *fyuA*, *cnf1*, *papA* (papA_F43 or papA_F16), *papC*, *hra* (encoding heat-resistant agglutinin), *irp2*, *chuA* (encoding outer membrane hemin receptor), *kpsE* (encoding capsule polysaccharide export inner-membrane protein) with the position of *fyuA* and *irp2* (both in the yersiniabactin siderophore cluster) relative to the other genes being the most variable. These isolates were associated with the kpsMIII_K96 group 3 capsule allele. The one isolate (P6_NW1_52/19) carrying *bla*_OXA-48_ carried the gene in an IncL/M plasmid, as would be expected.

The set included 3 representatives of ST73 from 2 different hospitals, which had completely different SNP addresses from one another (>250 SNP differences) showing that they were not epidemiologically related. These carried kpsMII_K5 (coding for polysialic acid transport protein; Group 2 capsule) but were otherwise highly similar in virulence gene content to the ST12 isolates. Similarly, they carried a section carrying *sitA*, *iss*, *iroN*, *focG*, *focCsfaE*, *mcmA*, *mchF*, *mchC*, *mchB* genes (in that order) followed by a section carrying *ompT*, *vat*, *pic* and *usp*. All also carried *chuA*, *gad*, *tcpC*, *irp2*, *fyuA*, *clbB*, *yfcV*, *terC*, *hra*, *cnf1* and *kpsE*. The *cnf1* gene was in a cluster with the *hylABCD* genes.

Indeed, most of the isolates sequenced carried *cnf1* and *hylBCD*, which were in a chromosomal genomic island (Fig. 3 a and b). Although these elements were carried by most isolates, the region was nevertheless quite mosaic in nature, with some isolates also carrying F1 and S fimbriae genes in this region, while others carried *pap* genes. Most of the virulence genes sought (e.g. the microcin (Fig. 3d) and colibactin (Fig. 3e) genes) were carried in such chromosomal islands, with clear differences in each region between isolates (Fig. 3).

## Discussion

While all the virulence factors are important, many were very common or very rare among our set, and we have chosen to concentrate on those for which there was evidence of a greater incidence among clinical isolates compared to that from screening isolates. Of course, colonisation is the first step in the evolution from commensal to pathogen causing extraintestinal infection, so not surprisingly, these factors were by no means confined to those from infections. In agreement with previously published work, it was clear from the long-read sequencing that the virulence factors were largely associated with PAIs on the chromosome, which have been linked with the emergence of virulence [17]. It was also clear that some sequence types, particularly STs 12, 73, 998 and 127, were strongly associated with multiple virulence factors, no matter whether they were from screening swabs or from blood or other infection sites. This was a consistent finding, not just within the study period, but also subsequently. Alhashash et al [25] noted an increase in incidence of representatives of ST73 causing clinical infections, particularly bacteraemias, and showed that, whilst they were relatively diverse, they mostly shared the same complement of virulence genes, a finding that was mirrored in this study. Similarly, STs 12, 80 and 998 were each associated with a consistent set of virulence genes, despite coming from many different hospitals and often having clearly distinct SNP addresses (at least 100 SNPs apart); STs 80 and 998 were notably strongly associated with the K1 capsule. It does seem that the number of virulence factors (VFs) is important, representing a continuum of acquisition in the evolution of the organism from harmless commensal to one capable of causing extraintestinal infections. Notably, most (99/120) representatives of ST131 carried relatively few VFs (mostly *malX* or *malX*, *sat* or *malX*, *papC*, *sat* among those sought), despite being considered one of the most important ExPEC lineages, highlighting the need to be aware of less well recognised linages. However, some representatives also carried *cnf1* and *hylBCD* (17), or *ibeA* (2). Worryingly, almost 9 % (53/593) of isolates in our study carried more than six virulence genes/gene sets, highlighting the potential for significant numbers of carriage isolates to cause extraintestinal infections. Of these, 13.6 % of isolates from neonatal screens carried 8 or more of the VFs sought, compared with 3.3 % of other screens. While this was in part associated with outbreaks in two hospitals, it is nevertheless concerning, especially considering that neonatal meningitis due to *E. coli* carries a high mortality (10%) and morbidity (30%) rate [26]. NMEC are characterised by K1 capsular antigens or the *ibeA* invasion gene, features particularly noted in STs 1193, 10, 998, 538, 80 and 141 in this study. However, these STs were not confined to isolates from neonates.

This work has provided a framework in which to assess isolates for virulence amongst a bewildering array of candidate genes and has highlighted types that harbour the greatest number of these virulence factors. It has also shown that a significant proportion of isolates carry numerous virulence factors with consequent increased potential to cause infection. These virulence factors are largely found in genomic islands, mosaic integrative structures that can be acquired by horizontal transfer.

## Conclusions

- *cnf1* (coding for cytotoxic necrotising factor), *clbK* (coding for colibactin), *focCDF* (coding for F1 fimbriae), *kpsM*_K1, *neuBD* (associated with the K1 capsule), *mchBC*, *mcmA* (coding for microcins), *papG*_alleleIII (part of a cluster encoding P fimbriae), *pic* (protein involved in intestinal colonization), *sfaE/sfafoCDE* (coding for S fimbriae), *tcpC* (encoding TLR domain containing-protein C) and *vat* (encoding toxin vacuolating autotransporter) were more prevalent among isolates from invasive infections than among those from carriage
- Representatives of STs 12, 73, 998 and 127 carried multiple of these virulence factors, no matter whether they were from screening swabs or from blood or other infection sites.
- Genes associated with the K1 capsule (*ibeA*, *neuBD*, *kpsM*_K1) were particularly found in STs 1193, 10, 998, 538, 80 and 141
- 9 % of isolates in our study carried more than six of these virulence genes/gene sets, highlighting the potential for significant numbers of carriage isolates to cause extraintestinal infections
- Isolates carrying multiple virulence factors were more prevalent from neonatal screens than those from general screens
- Most of the virulence factors were in chromosomal genomic islands, which are mosaic in nature

## Acknowledgements

This work is an analysis of sequences generated by a service provided by the Opportunistic Pathogens Section of AMRHAI, the Gastrointestinal Bacteria Reference Unit, the Central Sequencing Laboratory and the Bioinformatics Unit at UKHSA, Colindale and thanks are due to every member of all of those teams involved in this work. Thanks are also due to Jack Turton for analysis of nanopore sequences. We thank staff from hospitals for sending these isolates to us.

