## Supplemental Table S1 for "Virulence factors among isolates of extraintestinal *Escherichia coli* (ExPEC) from hospitals in the United Kingdom"

| Isolate | Sequence type | O:H type | Isolation site (category) | Other information | Virulence genes | Carbapenemase detected | Phylogenetic group  (*chuA*,TspE4C2, *yjaA*) |
| --- | --- | --- | --- | --- | --- | --- | --- |
| P1_SE1_24/20 | 80 | O75:H7 | Urine (U) | Mother of N1 | *cblK,cnf1,focCDF,hylBCD,ibeA,mchBC,neuBD,sfaE, KpsM*_K1*,malX,mcmA,pic,vat* | - | B2 (+,-,+) |
| N1_SE1_24/20 | 80 | O75:H7 | CSF (CSF) | Baby of P1 | *cblK,cnf1,focCDF,hylBCD,ibeA,mchBC,neuBD,sfaE, KpsM*_K1*,malX,mcmA,pic,vat* | - | B2 (+,-,+) |
| N17_EE2_40/20 | 80 | O7:H7 | Blood | Requested detection of K1 | *clbK,cnf1,focCDF,hylCD,mchBC,ibeA,neuBD,sfaE, malX,mcmA,papC,pic,vat,KpsM*_K1 | - | B2 (+,+,+) |
| P26_Y&H2_02/21 | 80 | O75:H7 | Sputum | From 1 month old baby | *clbK,cnf1,focCDF,hylBCD,ibeA,kpsM*_K1*,mchBC,neuBD,sfaE, malX,mcmA,pic,vat* | - | B2 (+,+,+) |
| N2_WM1_50/19 | 998 | O50O2:H6 | pooled swabs (NS) | Same strain as P2, N4 | *cblK,cnf1,focCDF,hylBCD,ibeA,sfaE,tcpC, malX,vat* | - | B2 (+,+,+) |
| P2_WM1_52/19 | 998 | O50O2:H6 | infection screen (AS) | Same strain as N2, N4 | *clbK,cnf1*,focCDF,hylBCD,ibeA,sfaE,tcpC, malX,vat* | - | B2 (+,+,+) |
| N3_WM1_26/19 | 998 | O50O2:H6 | pooled swabs (NS) | Distinct from other ST998 by PFGE | *clbK,cnf1,focCDF,hylBCD,ibeA,neuBD,sfaE,tcpC, KpsM*_K1*,malX,vat* | - | B2 (+,+,+) |
| N4_WM1_52/19 | 998 | O50O2:H6 | Eye (other) | same strain as N2, P2 | *clbK,cnf1,focCDF,hylBCD,ibeA,sfaE,tcpC, malX,pic,vat* | - | B2 (+,+,+) |
| P19_NW1_44/20 | 998 | O50O2:H6 | Rectal screen swab | Screening for carbapenemases | *clbK,cnf1,focCDF,hylBCD,ibeA,KpsM*_K1*,neuB,neuD,sfaE,tcpC. malX,vat* | OXA-181 | B2 (+,+,+) |
| P25_NW1_ 08/21 | 998 | O50O2:H6 | Rectal screen |  | *clbK,cnf1,focCDF,hylBCD,ibeA,KpsM*_K1*,neuB,neuD,sfaE,tcpC, malX,vat* | OXA-181 | B2 (+,+,+) |
| P24_SE2_ 52/20 | P3595 (13,52,10,14,*17,25,17)  SLV of 998 | O50O2:H6 | Blood | From 3 month old baby | *clbK,cnf1,focCDF,hylBCD,mchBC,mcmA,sfaE,tcpC, malX,papC,papG*_alleleIII*,tsh,vat* | - | B2 (+,+,+) |
| P3_L1-52/19 | 625 | O6:H7 | Rectal (AS) | screen | *clbK,cnf1,focCDF,hylBCD,ibeA,mchBC,sfaE,tcpC, malX, mcmA, papC, papG*_alleleIII*,vat* | - | B2 (+,+,+) |
| P4_EM1_41/19 | 141 | O50O2:H6 | CSF (CSF) | None given | *clbK,cnf1,focCDF,hylBCD,mchBC,neuBD,sfaE,tcpC, KpsM*_K1*,malX,mcmA,vat* | - | B2 (+,+,+) |
| P5_EM1_41/19 | 12 | O4:H5 | CSF (CSF) | None given | *clbK,cnf1,focCDF,hylBCD,mchBC,sfaE,tcpC, malX,mcmA,papC,papG*_alleleIII*,vat* | - | B2 (+,-,+) |
| P6_NW1_52/19 | 12 | O4:H5 | rectal screen (AS) | screen | *clbK,cnf1,focCDF,hylBCD,mchBC,sfaE,tcpC, malX,mcmA,papC,papG*_alleleIII*,vat* | OXA-48 | B2 (+,-,+) |
| P7_NW2_18/20 | 12 | O4:H5 | Blood (B) | Sepsis, stillbirth | *clbK,cnf1,focCDF,hylBCD,mchBC,sfaE,tcpC, malX,mcmA,papC,papG*_alleleIII*,vat* | - | B2 (+,-,+) |
| N9_EE1_25/20 | 12 | O4:H5 | Faeces (NS) | None given | *cnf1,focCDF,hylBCD,mchBC,sfaE,tcpC, malX,mcmA,papC,papG*_alleleIII*,vat* | - | B2 (+,-,+) |
| N16_EE1_ 42/20 | 12 | O4:H5 | faeces | neonate | *clbK,cnf1,focCDF,hylBCD,mchBC,sfaE,tcpC, malX,mcmA,papC,papG*_alleleIII*,vat* | - | B2 (+,+,+) |
| N18_L5_ 48/20 | 12 | O4:H5 | Rectal swab | neonate | *clbK,cnf1,hylBCD, tcpC, malX,papC, papG*_alleleIII*,vat* | - | B2 (+,+,+) |
| P22_SW1_51/20 | 12 | O4:H5 | Pus-neck abscess |  | *clbK,cnf1,focCDF,hylBCD,mchBC,sfaE,tcpC, malX,mcmA,papC,papG*_alleleIII*,sat,vat* | - | B2 (+,+,+) |
| N21_EE1_52/20 | 12 | O4:H5 | faeces | Routine screen | *clbK,cnf1,focCDF,hylBCD,mchBC,sfaE,tcpC, malX,mcmA,papC,papG*_alleleIII*,vat* | - | B2 (+,+,+) |
| N5_WM1_04/20 | 73 | O6:H1 | line tip (other) | None given | *clbK,cnf1,focCDF,hylBCD,mchBC,sfaE,tcpC, malX,mcmA,pic,sat,upaH,vat* | - | B2 (+,+,+) |
| P14_Y&H1_29/20 | 73 | O22:H1 | Blood |  | *clbK,cnf1,focCDF,hylBCD,mchBC,sfaE, malX,mcmA,papC,papG*_alleleIII*,pic,upaH,vat** | - | B2 (+,+,+) |
| P15_Y&H1_29/20 | 73 | O6:H1 | blood |  | *clbK,cnf1,focCDF,hylBCD,mchBC,sfaE,tcpC, malX,mcmA,papC,pic,sat,upaH*,vat* | - | B2 (+,+,+) |
| N11_WM1_31/20 | 73 | O6:H1 | Pooled swabs | 5-SNP cluster with N12, N13 and N14 | *clbK,cnf1,focCDF,hylBCD,mchBC,sfaE,tcpC, malX, mcmA, pic, upaH, vat* | - | B2 (+,+,+) |
| N12_WM1_31/20 | 73 | O6:H1 | Nasopharyngeal aspirate | 5-SNP cluster with N11, N13 and N14 | *clbK,cnf1,focCDF,hylBCD,mchBC,sfaE,tcpC, malX,mcmA,pic,upaH,vat* | - | B2 (+,+,+) |
| N13_WM1_31/20 | 73 | O6:H1 | Pooled swabs | 5-SNP cluster with N11, N12 and N14 | *clbK,cnf1,focCDF,hylBCD,mchBC,sfaE,tcpC, malX,mcmA,pic,upaH,vat* | - | B2 (+,+,+) |
| N14_WM1_31/20 | 73 | O6:H1 | Pooled swabs | 5-SNP cluster with N11, N12 and N13 | *clbK,cnf1,focCDF,hylBCD,mchBC,sfaE,tcpC, malX,mcmA,pic,upaH,vat* | - | B2 (+,+,+) |
| P16_WM2_33/20 | 73 | O6:H1 | blood |  | *clbK,focCDF,mchBC,sfaE,tcpC, malX,mcmA,pic,sat,upaH,vat* | - | B2 (+,+,+) |
| P18_SW2_41/20 | 73 | O6:H1 | Blood | ESBL | *clbK,cnf1,focCDF,hylBCD,mchBC,sfaE,tcpC, malX,mcmA,papC,pic,sat,upaH,vat* | - | B2 (+,+,+) |
| N22_Y&H2_02/21 | 73 | O6:H1 | Sputum | Neonate | *clbK,cnf1,focCDF,hylBCD,mchBC,sfaE,tcpC, malX,mcmA,pic,sat,upaH*,vat* | - | B2 (+,+,+) |
| P27_WM1_09/21 | 73 | O6:H1 | Nose and groin |  | *cblK,cnf1,focCDF,hylBCD,mchBC,sfaE,tcpC, malX,mcmA,papC,pic,sat,upaH,vat* | - | B2 (+,+,+) |
| N23_Y&H2_09/21 | 73 | O18:H1 | ET Tube tip | Neonate | *cnf1,focCDF,hylBCD,mchBC,sfaE,tcpC, malX,mcmA,pic,papG*_alleleIII*,sat,upaH,vat* | - | B2 (+,+,+) |
| E_S1_10/20 | P6 (*36,24,9,13,17,11,25) (SLV of ST73) | O22:H1 | Environmental (E) | From mascerator | *clbK,cnf1,focCDF,hylBCD,mchBC,sfaE, malX,mcmA,pic,upaH,vat* | - | B2 (+,+,+) |
| N15_EE2_39/20 | 2013 (CC73) | O6:H1 | Blood | septic | *clbK,cnf1,focCDF,hylBCD,mchBC,sfaE, malX,mcmA,papC,papG*_alleleIII*,pic,upaH,vat* | - | B2 (+,+,+) |
| P8_SW1_27/20 | 141 | O50O2:H6 | Thigh tissue (other) | Abscess, septic arthritis | *clbK,cnf1,focCDF,hylBCD,neuBD,sfaE,tcpC, KpsM*_K1*,malX,papC,papG*_alleleIII*,sepA,vat,virF* | - | B2 (+,+,+) |
| P9_NW1_07/20 | 127 | O6:H31 | rectal screen (AS) | Elderly, screen | *clbK,cnf1,focCDF,hylBCD,sfaE, malX,pic,vat* | OXA-48 | B2 (+,+,+) |
| P10_EM1_41/19 | 127 | O6:H31 | CSF (CSF) | CSF | *clbK,cnf1,focCDF,hylBCD,sfaE,tcpC, KpsM*_K1*,malX,mcmA,papC,papG*_alleleIII*,vat* | - | B2 (+,+,+) |
| P11_NE1_45/19 | 127 | O6:H31 | kidney urine (U) | Not given | *clbK,cnf1,focCDF,hylBCD,sfaE,tcpC, malX,mcmA,papC,vat* | OXA-48 | B2 (+,+,+) |
| P17_W1_33/20 | 127 | O6:H31 | Pus | Abscess | *clbK,cnf1,focCDF,hylBCD,sfaE,tcpC, malX,mcmA,papC,vat* | - | B2 (+,+,+) |
| N19_Y&H2_45/20 | 127 | O6:H31 | endotracheal tube tip | neonate | *clbK,cnf1,focCDF,hylBCD,sfaE,tcpC, malX,mcmA,papC,papG*_alleleIII*,vat* | - | B2 (+,+,+) |
| P20_NW1_48/20 | 127 | O6:H31 | Rectal swab | Stroke unit | *clbK,cnf1,focCDF,hylBCD,sfaE, malX,vat* | OXA-48 | B2 (+,+,+) |
| N6_S1_40/19 | 416 (CC95) | O18ac:H7 | Blood (B) | 5 SNP cluster with N7 | *clbK,focCDF,ibeA,neuBD,sfaE, KpsM*_K1*,malX,vat* |  | B2 (+,+,+) |
| N7_S1_41/19 | 416 (CC95) | O18ac:H7 | CSF (CSF) | 5 SNP cluster with N6 | *clbK,focCDF,ibeA,neuBD,sfaE, KpsM*_K1*,malX,vat* |  | B2 (+,+,+) |
| N8_L2_07/20 | 1859 | O99:H6 | Blood (B) | Sepsis | *cnf1,focCDF,hylBCD,ibeA,mchBC,neuBD,sfaE, KpsM*_K1*,malX,mcmA,pic,vat* | - | B2 (+,+,+) |
| P12_NW1_38/19 | 372 | O83:H31 | CSU (U) | urine | *cnf1,focCDF,hylBCD,ibeA,mchBC,sfaE, malX,mcmA,papC,papG*_alleleIII*,vat* | OXA-48 | B2 (+,+,+) |
| N10_L3_26/20 | 4456 | O83:H4 | blood culture (B) | Meningitis | *focCDF,ibeA,mchBC,neuBD,sfaE, KpsM*_K1*,malX,mcmA,papC,papG,vat* | - | B2 (+,-,+) |
| P13_Y&H1_29/20 | P3552 (36,24,550,*13,17,11,25) | O6:H1 | Blood |  | *cnf1,hylBCD,tcpC, malX,papC,papG,pic,sat,upaH,vat* | - | B2 (+,+,+) |
| P21_NW1_ 48/20 | 544 | O4:H5 | Rectal screen |  | *clbK,cnf1,focCDF,hylBCD,mchBC,sfaE,tcpC, malX,mcmA,papC* | - | B2 (+,-,+) |
| N20_EE1_50/20 | 83 | O6:H5 | faeces | Weekly routine testing | *clbK,cnf1,focCDF,hylBCD,ibeA,mchBC,mcmA,sfaE, malX,mcmA,papC,papG*_alleleIII*,vat* | - | B2 (+,+,+) |
| P23_NW1_52/20 | 681 | O8:H10 | Rectal screen swab | From elderly patient | *clbK,focCDF,mchBC,mcmA,sfaE, malX,mcmA,pic,upaH* | OXA-48 | B2 (+,+,+) |
| N24_Y&H2_10/21 | 2017 | O22:H14 | blood | Neonate | *cnf1,focCDF,hylBCD,ibeA,mchBC,sfaE, malX,mcmA,vat* | - | B2 (+,+,+) |
| P28_WM1_12/21 | 2015 | O50O2:H14 | Groin |  | *clbK,cnf1,hylBCD,ibeA,mchBC,sfaE, malX,mcmA,papC,sat,papG*_alleleIII*,vat* | - | B2 (+,+,+) |

Table S1. Details of the 53 isolates carrying more than 6 of the virulence gene/gene sets sought. Isolates were labelled by patient (P1-P28, N1-N24, where those beginning with N are from neonates), hospital (by region and number within that region) and date of isolation (week/year). Regions were EE, East of England, EM, East Midlands, WM, West Midlands, L, London, NE, North East, NW, North West, SE, South East, SW, South West, S, Scotland, W, Wales, Y&H, Yorkshire and Humber. There was one environmental isolate (E).
